# Regulation of sociality and neuronal development by the translation initiation factor *eIF4H1* in *Drosophila*

**DOI:** 10.64898/2026.11.06.764618

**Authors:** Erik Nolan, Iris Chin, Cassondra Vernier, Yehuda Ben-Shahar

**Affiliations:** Department of Biology, Washington University in St. Louis, St. Louis, MO, USA; Gladstone Institute of Neurological Disease, Gladstone Institutes, San Francisco, CA, USA; Biology Department, University of Wisconsin - Stout Polytechnic, WI, USA

## Abstract

Most animal species exhibit some level of sociality. However, whether sociality-related traits evolved independently, or are instead derived from common core mechanisms, remains controversial. Nevertheless, an emerging theoretical framework stipulates that key aspects of how animals interact with their social environments may be regulated by a phylogenetically conserved genetic toolkit. Therefore, we hypothesized that mutations that lead to social deficits in specific human disorders may implicate genes that play conserved roles in the regulation of core social traits across Animalia. Specifically, here we focus on genes associated with Williams syndrome (WS), a rare human multi-system disorder characterized in part by hypersocial cognitive traits. WS is caused by a hemizygous chromosomal microdeletion of 25-28 protein-coding genes. Consequently, it has been hypothesized that haploinsufficiency in at least one of these genes plays an important role in the development and/or function of social neural circuits in humans. Because the *Drosophila* genome encodes orthologs of 17 of the 28 genes deleted in WS, we reasoned that at least some of them may serve a conserved role in the regulation of social traits across flies and humans, as predicted by the genetic toolkit framework. Using a neuron-specific RNAi knockdown screen, we identified *eIF4H1*, an ortholog of a highly conserved subunit of the eukaryotic translation initiation complex eIF4F, as a candidate gene for neuronal regulation of sociality in the fly. We show that *eIF4H1* expression is predominantly neuronal in the adult fly brain, and that its action plays a differential role in the organization of specific dopaminergic neuron subtypes that are associated with sociality and cognitive functions. These results indicate that *eIF4H1* may serve a conserved function in regulating the development of the neuronal circuits that regulate social behaviors, thus supporting the idea of a conserved sociality genetic toolkit.

## Introduction

Animal sociality, which spans a continuum from transient aggregations to highly organized eusocial systems, is thought to have evolved multiple independent times across the animal phylogeny (Alexander, 1974; Nowak et al., 2010). Despite this seemingly convergent evolution, growing evidence suggests that social traits across distant taxa are often driven by shared molecular and cellular mechanisms (Robinson et al., 2008; O’Connell & Hofmann, 2011; Ben-Shahar et al., 2002; Lu et al., 2014). Core neuromodulatory systems such as the biogenic amines dopamine and serotonin, diverse neuropeptides, and hormones recurrently regulate social interactions, motivation, and group cohesion across taxa as distant as insects and vertebrates (Donaldson & Young, 2008; Robinson et al., 2019). At the cellular level, conserved neural circuits and gene regulatory networks, including those governing sensory processing, reward, and developmental plasticity, are repeatedly co-opted to produce social phenotypes (O’Connell & Hofmann, 2011). These findings suggest that the evolution of social behaviors may be driven by the repeated recruitment of conserved neurobiological pathways, rather than frequent emergence of entirely new molecular and cellular mechanisms across independent taxa. However, the specific identities of the core gene networks that drive animal sociality, and the mechanisms by which they exert their functions, remain a mystery for most species.

A potentially productive approach to understanding sociality at the molecular and cellular levels can be derived from many observations which indicate that analogous complex phenotypes are often associated with the conserved action of specific genes and signaling pathways. Evolutionary studies of embryonic development across distant metazoan taxa uncovered several highly conserved genetic pathways, such as the Hox gene family (McGinnis & Krumlauf 1992; Carroll, 1995). These observations were subsequently formalized by the evolutionary “genetic toolkit” framework. This framework stipulates that morphological diversity often arises on the evolutionary timescale not via novel genes, but from variation in when, where, and how conserved developmental “toolkit genes” exert their functions (Carroll, 2000). More recently, discoveries about the deep conservation of biological pathways that underlie neuronal differentiation, circuit formation, and postmitotic neurophysiology across distant animal taxa suggest that this genetic toolkit concept may be applicable to studies of neural functions and behaviors (Rittschof & Robinson, 2016; Toth & Robinson, 2007). Consequently, identifying a conserved “sociality genetic toolkit” would imply that even some of the most complex social traits exhibited by different animal species are driven by conserved sets of genes that regulate relevant processes such as reproduction, response to stressors, and the reward pathways (Bloch & Grozinger, 2011; Robinson et al., 2008)

Human genetic disorders associated with abnormal sociocognitive profiles can provide experimental opportunities for testing mechanistic hypotheses within the sociality genetic toolkit conceptual framework. One example is Williams syndrome (WS), a rare multi-systemic disorder caused by a hemizygous microdeletion of 25–28 protein-coding genes at chromosome region 7q11.23 (Kozel et al., 2021). The syndrome is characterized by highly stereotypic cardiovascular abnormalities, distinctive craniofacial features, and a striking cognitive-behavioral profile that includes hypersociability, heightened social approach, strong empathy, and reduced social fear, often accompanied by increased anxiety and atypical sensory processing (Järvinen et al., 2013). Neurobiologically, individuals with WS show alterations in circuits involving the amygdala, prefrontal cortex, and reward pathways, along with dysregulation of neuromodulatory systems such as oxytocin and dopamine that are central to social motivation and salience (Barak & Feng, 2016). Consequently, the combination of the molecularly well-defined chromosomal deletion of a small number of protein-coding genes, the molecular and biochemical conservation of the affected genes across Animalia, and the high syndromic phenotypic penetrance in individuals with WS indicate that it could serve as a powerful hypothesis-generating tool for gaining mechanistic insights into potentially conserved genetic and neuronal features of animal sociality.

Therefore, here we tested the hypothesis that at least some of the genes covered by the Williams syndrome microdeletion in humans are also important nodes in the gene networks that drive social traits in the vinegar fly *Drosophila melanogaster*. Our initial gene homology analysis revealed that the *Drosophila* genome encodes orthologs for at least 17 of the human protein coding genes covered by the WS deletion. Using a neuronal RNAi knockdown screen for group sociality (Simon et al., 2012; Couzin, 2009), we found that the knockdown of at least one of these orthologs, the highly conserved eukaryotic translation initiation factor *eIF4H1* (Marintchev et al., 2009; Rogers et al., 2001), leads to abnormal social interactions in adult *Drosophila*. Subsequently, analysis of males with either neuron-specific RNAi knockdown or a knockout allele of *eIF4H1* revealed that this gene plays a role in aspects of social-and reproduction-related behaviors in male flies. High resolution imaging of a GFP-tagged *eIF4H1* allele indicate that eIF4H1 expression is broadly enriched in postmitotic neurons of the fly brain, and that the impact of its null allele on the differentiation and morphology of specific neuromodulatory dopaminergic subpopulations that drive reproductive behaviors and various cognitive functions is variable. Lastly, we show that loss of *eIF4H1* results in aberrant dendritic morphology in peripheral sensory neurons. Together, our data suggest that *eIF4H* is a candidate component of a conserved genetic toolkit that is responsible for driving some of the core aspects of animal sociality.

## Results

### *Genetic screen of Williams syndrome orthologs in Drosophila reveals eIF4H1 as a* regulator of sociality

To test the hypothesis that at least one of the Williams syndrome-associated genes may play a conserved role in regulating animal social behavior, we first searched for *Drosophila* orthologs of the human genes deleted in WS by utilizing the DRSC Integrative Ortholog Prediction Tool (DIOPT) (Hu et al., 2011). DIOPT analysis identified at least 17 *Drosophila* genes that encode likely orthologs of protein coding genes deleted in WS (**Table 1**). Next, we utilized the TRiP VALIUM20 UAS-RNAi transgenic line collection (Perkins et al., 2015) to knockdown eight of these orthologs in the fly nervous system by using the pan-neuronal *elav*-GAL4 driver (Lin & Goodman, 1994). For the human gene *FZD9*, the DIOPT analysis identified both *fz3* and *fz4* as likely orthologs (**Table 1**). When available, multiple VALIUM20 UAS-RNAi lines that target the same *Drosophila* gene were used. We found that pan-neuronal knockdown of two genes – *Syx1A* and *CG10903* – resulted in F1 lethality (**Table 1**). Subsequently, viable F1 adults were used to investigate the impact of RNAi gene knockdowns on fly social behavior by using the social space displacement paradigm (Simon, 2012) (**Fig 1A**). We found that the neuronal knockdown of *eIF4H1* was the only one to impact behavior in the social space assay (**Fig 1B-D**). Using a climbing assay (Lu et al., 2012; Zheng et al., 2014), we found that this effect was not due to general locomotion deficits (**Fig 1E**). We also investigated whether the effect of neuronal *eIF4H1* knockdown on sociality was specific to the social space paradigm by studying its effects on stereotypic male courtship behaviors (McKinney et al., 2026; Vernier et al., 2023; Villella & Hall, 2008; Crossley & Zuill, 1970; Sokolowski, 2001). We found that the pan-neuronal knockdown of *eIF4H1* in males increased their latency to courtship initiation (**Fig 1F**). In contrast, proportion of total time spent courting (courtship index) was unaffected (**Fig 1G**). Together, these data indicate that *eIF4H1* action is necessary for the development and/or post-mitotic functions of sociality-relevant neuronal circuits in *Drosophila*.

**Figure 1.**
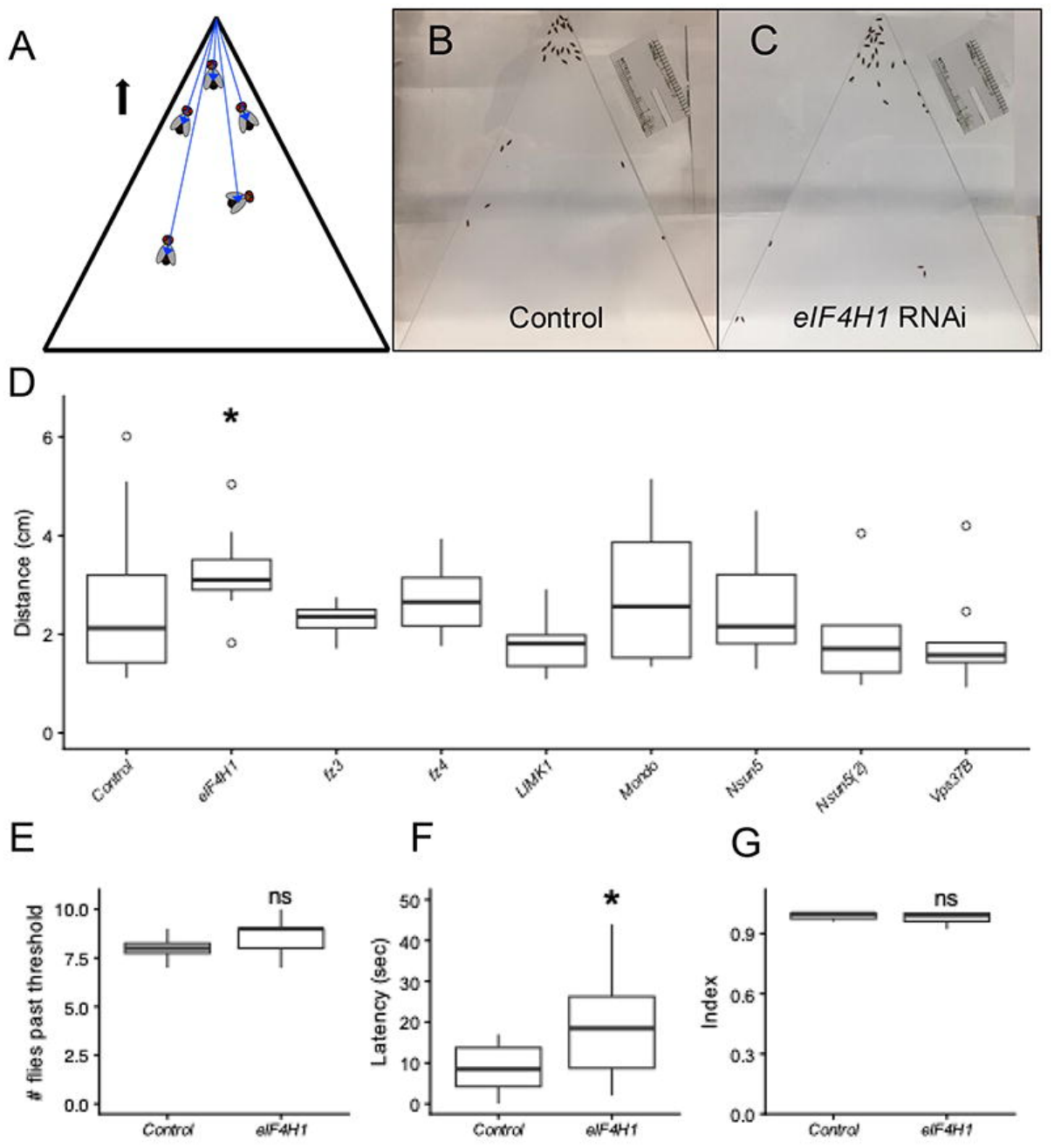
Neuronal RNAi knockdown screen reveals *eIF4H1* as a regulator of fly social behavior. A) Diagram of social space arena, arrow indicates geotactic preference of flies to move up to the top of the arena. Blue lines indicate distance to top of arena that is averaged across all flies in a trial. B & C) Representative social space assay images of control and *eIF4H1-*RNAi lines. D) Average group distances to arena top for trials of gene RNAi knockdowns. Sample sizes: control n = 29, *eIF4H1-*RNAi n = 11, all other RNAi lines n > 8. * indicates *p*-value < 0.05 before multiple comparison adjustment (Adj. *p*-value = 0.65, Dunn’s test following Kruskal-Wallis Rank Sum). E) Number of flies able to climb past 10 cm threshold in locomotor control experiment (n = 10, two-tailed t-test, ns = not significant) F) Courtship latency and G) index of *eIF4H1*-RNAi male flies (n =16-19, Wilcoxon rank sum test, * *p*-value < 0.05, ns = not significant). In all boxplots shown, bottom and top of the box represent the 25th and 75th percentiles (interquartile range, IQR). The central line indicates the median, whiskers extend to 1.5 times the IQR / minimum and maximum values, and outliers are plotted as individual points.

**Table 1.** Drosophila Williams syndrome gene orthologs. DIOPT score and rank reflect strength of identified fly gene as ortholog of respective human Williams-associated gene. Gray boxes indicate genes associated with Williams syndrome with no detectable fly orthologs. Screened genes are in bold. Red indicates fly orthologs that were lethal upon neuronal knockdown.

|  | Human Gene | Fly Orthologs | DIOPT Score | Rank |
|---|---|---|---|---|
| 1 | POM121 | *CG14712* | 2 | High |
| 2 | **NSUN5** | *Nsun5* | 13 | High |
| 3 | TRIM50 |  |  |  |
| 4 | FKBP6 | *shu* | 13 | High |
| 5 | **FZD9** | *fz3* | 5 | Moderate |
|  |  | *fz4* | 4 | Moderate |
| 6 | BAZ1B | *tou* | 2 | Moderate |
| 7 | BCL7B | *BCL7-like* | 11 | Moderate |
| 8 | TBL2 |  |  |  |
| 9 | **MLXIPL** | *Mondo* | 12 | High |
| 10 | DNAJC30 | *CG11035* | 10 | High |
| 11 | BUD23 | *CG10903* | 12 | High |
| 12 | **VPS37D** | *Vps37B* | 9 | Moderate |
| 13 | **STX1A** | *Syx1A* | 13 | High |
| 14 | ABHD11 | *CG2059* | 13 | High |
| 15 | CLDN3 |  |  |  |
| 16 | CLDN4 |  |  |  |
| 17 | **METTL27** | *CG10903* | 2 | Moderate |
| 18 | TMEM270 |  |  |  |
| 19 | ELN |  |  |  |
| 20 | **LIMK1** | *LIMK1* | 11 | High |
| 21 | **EIF4H** | *eIF4H1* | 11 | High |
| 22 | LAT2 |  |  |  |
| 23 | RFC2 | *RfC4* | 13 | High |
| 24 | CLIP2 | *CLIP-190* | 8 | Moderate |
| 25 | GTF2IRD1 |  |  |  |
| 26 | GTF2I |  |  |  |
| 27 | NCF1 |  |  |  |
| 28 | GTF2IRD2 |  |  |  |

### Endogenous eIF4H1 protein is expressed in neurons of the adult central brain

The *eIF4H1* gene encodes a component of the eIF4F eukaryotic translation initiation complex, which is conserved from yeast to humans (Sonenberg & Hinnebusch, 2009; Hinnebusch, 2014) (**Fig 2A**). The eIF4F complex is responsible for the initial and rate-limiting steps of translation, including the recognition and binding of the 5’ m⁷G cap, followed by processing of the 5’ untranslated region (UTR) and recruitment of downstream translation machinery (Gingras et al., 1999). Within the eIF4F complex, the mutually exclusive binding of eIF4H or its paralog eIF4B is thought to augment the RNA helicase activity of eIF4A, which contributes to the initiation of translation by resolving secondary structures in mRNAs with long and/or complex 5’ UTRs (Rogers et al., 2001; Rozovsky et al., 2008; Chu et al., 2016; Marintchev et al., 2009). Extensive regulation of eIF4F complex activity has been shown to be important for the differentiation, development, and function of peripheral and central neurons in insect and mammalian species (Santini et al., 2013; Amorim et al., 2018; Costa-Mattioli et al., 2009; Rode et al., 2018). Therefore, based on the outcomes of our initial behavioral RNAi screen, we hypothesized that *eIF4H1*-dependent regulation of neuronal mRNA translation supports the developmental and/or physiological function of neural circuits that contribute to social behaviors.

**Figure 2.**
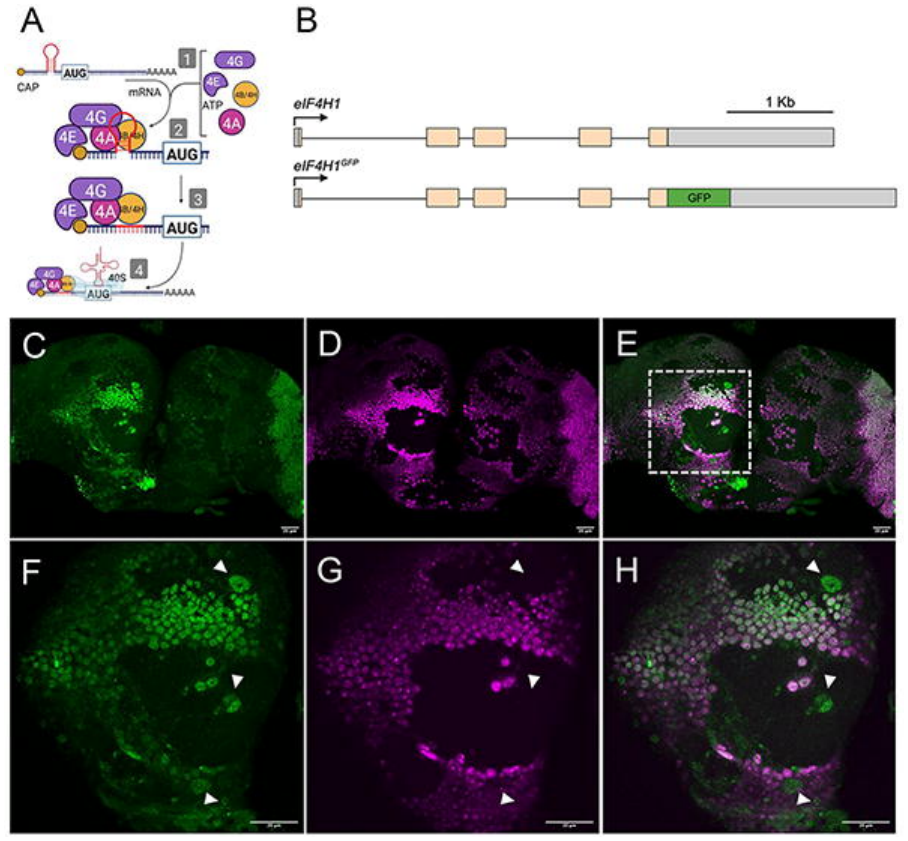
eIF4H1^GFP^ is expressed in neurons of the adult fly brain. A) Schematic of eIF4H function within the eIF4F complex; eIF4H and its paralog eIF4B bind to the RNA helicase subunit eIF4A to facilitate the initiation of translation for mRNAs with long and/or complex 5’ UTRs. B) Schematic of *eIF4H1^GFP^* allele (bottom) compared to the wild-type *eIF4H1* gene locus (top). C-H) Fluorescent microscopy images of a representative brain stained for eIF4H1^GFP^ (C) and ELAV (D), with merged channels shown in (E). Dashed box in (E) indicates area shown at higher magnification in (F-H). White arrowheads indicate non-neuronal (ELAV^-^) but eIF4H1-positive (GFP^+^) cells.

Following this overarching hypothesis, we first sought to define the cellular context within the nervous system in which *eIF4H1* could influence social behaviors. We therefore profiled its spatial expression pattern in the adult brain. First, we generated a GFP-tagged *eIF4H1* allele (*eIF4H1*^GFP^) by inserting the GFP coding sequence (CDS) in-frame at the gene’s 3’ CDS terminus using our previously published scarless CRISPR/Cas9-dependent genome editing approach (**Fig 2B**) (Bier et al., 2018; Hill et al., 2019). Next, we detected the distribution of the native eIF4H1 protein tag by using anti-GFP immunofluorescence and high-resolution confocal imaging (Hill et al., 2019; Vernier et al., 2023). To examine whether eIF4H1 is specifically expressed in postmitotic neurons, we co-stained adult brains with an anti-ELAV antibody, a general marker for postmitotic neurons (Robinow & White, 1991; Sanfilippo et al., 2016). Our imaging analysis revealed that the eIF4H1^GFP^ allele is broadly expressed in adult neurons across the central brain, as well as several large ELAV-negative cells of unknown function (**Fig 2C-H**). These data suggest a potential role for eIF4H1 in contributing to neuronal physiology.

### A loss-of-function eIF4H1^null^ allele is associated with abnormal sociality

As a part of our overarching hypothesis that *eIF4H1* activity regulates specific aspects of neuronal biology, we sought to establish cellular paradigms in which to further explore the activity of this gene. Consequently, we generated an *eIF4H1* null allele (*eIF4H1^null^*) by using CRISPR/Cas9 genome editing strategy to insert a 3xP3>DsRed cassette in the last coding exon of the *eIF4H1* locus (Bier et al., 2016; Hill et al., 2019) (**Fig 3A**). We found that while the X-linked *eIF4H1^null^* allele is homozygous-viable, it is associated with delayed development and strongly reduced viability and fecundity in eclosed hemizygous males **(Fig S1)**. The *eIF4H1^null^*stock therefore was therefore maintained as heterozygous over a wild type X chromosome in females.

**Figure 3.**
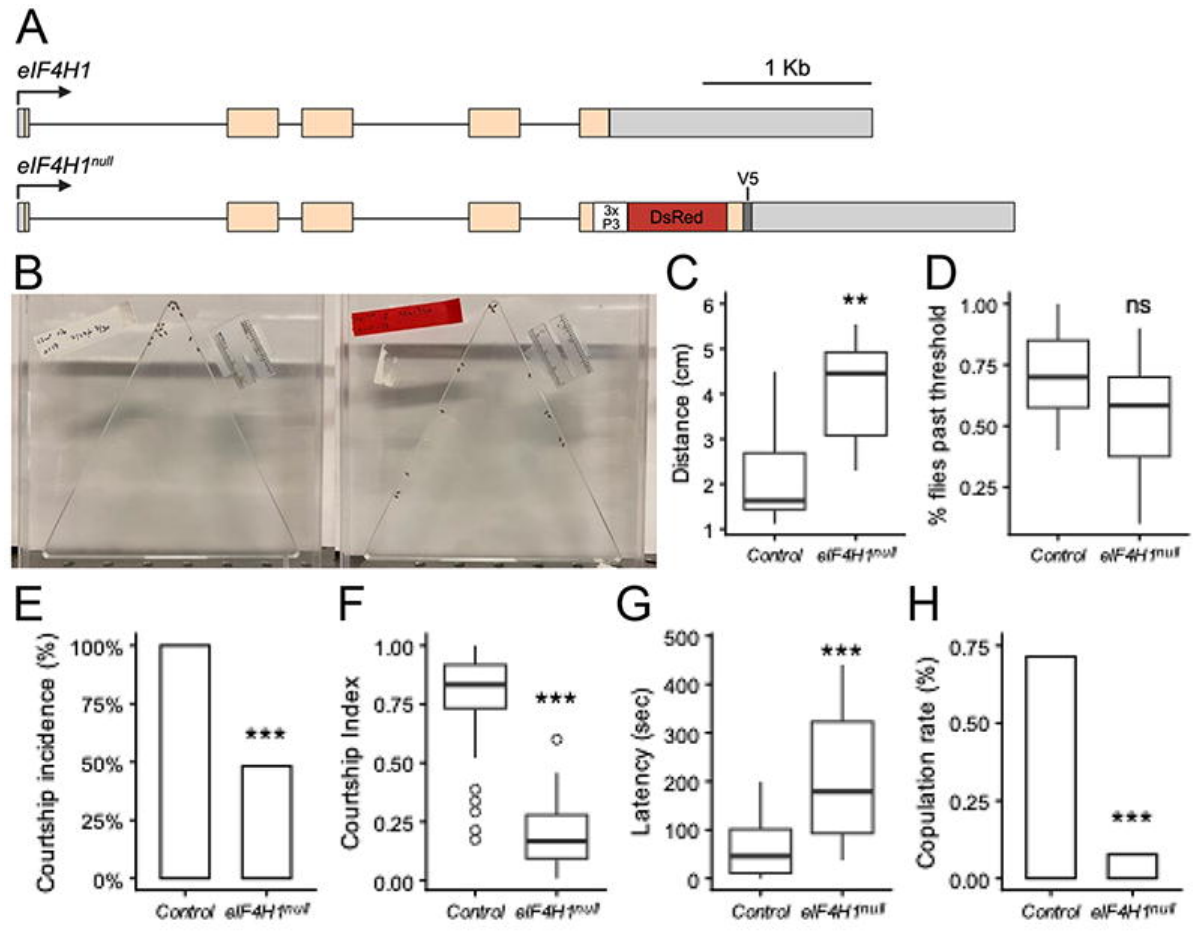
***eIF4H1^null^*results in impaired social behaviors.** A) Schematic of *eIF4H1^null^*allele, organized as in Fig 2. B) Representative images of social space assay results for Canton-S control (left) and *eIF4H1^null^* (right). C) Mean distances to top of arena for *eIF4H1^null^* social space trials (n = 8-10, Wilcoxon rank sum test). D) Number of flies able to climb past 10 cm threshold in locomotor control experiment (n = 10, *p*-value = 0.067, t-test). E-H) *eIF4H1^null^*courtship metrics, with n = 27-28 flies tested prior to exclusion of non-courting flies: courtship incidence (E), index (F), latency (G), and copulation rate (H). Courtship incidence and copulation rate were analyzed using Fisher’s exact test, while courtship index and latency were tested with Wilcoxon rank-sum test. * = *p*-value < 0.05, ** = *p*-value < 0.005, *** = *p*-value < 0.001. Box plot design as in Figure 1.

Next, we used the social space paradigm (**Fig 1A**) to show that, similarly to the neuron-specific RNAi knockdown (**Fig 1D**), the *eIF4H1^null^* allele is associated with an increase in the social space preference index (**Fig 3B-C**). This effect was not due to deficits in general motor skills, as *eIF4H1^null^* males performed similarly to wild type animals in a climbing assay (**Fig 3D**).

We also studied the courtship behavior of *eIF4H1^null^*males towards wild-type females. We found that *eIF4H1^null^* males exhibited reduced male courtship frequency, with only 50% of *eIF4H1^null^*males displaying any courtship during the trial (**Fig 3E**). Furthermore, the null males that did court exhibited longer latency to courtship onset (**Fig 3F**), decreased courtship index (**Fig 3G**), and a lower successful copulation rate (**Fig 3H**).

### *Differential effects of eIF4H1^null^ on dopaminergic neuron organization in the adult* brain

Dopaminergic neurons (DANs) play conserved roles in the central modulation of social behaviors and cognition across animals, including in *Drosophila* (Barron et al., 2010; Aragona et al., 2006; Gunaydin et al., 2014; Li et al., 2020; Zhang et al., 2016; Keleman 2012; Alekseyenko et al., 2013; Handler et al., 2019; Aso et al., 2010; Aso & Rubin, 2016). Furthermore, studies of WS and other syndromes associated with social deficits suggest that reduced social cognition abilities are driven, at least in part, by the abnormal development of specific elements within the dopaminergic system (Dichter et al., 2012; Kleberg et al., 2023; Mobbs et al., 2007). Similarly, a recent study of brain monamine levels in a mouse WS model revealed reduced dopaminergic signaling in multiple brain regions, further implying that impaired dopaminergic signaling may underlie some of abnormal cognitive and social symptoms associated with WS (Aman et al. 2025). We therefore hypothesized that *eIF4H1* activity might be necessary for the development of certain dopaminergic neurons in the *Drosophila* brain.

The *Drosophila* dopaminergic system is composed of several neuronal subtypes that are morphologically and functionally distinct (Kasture et al., 2018; Mao & Davis, 2009; White et al., 2010). These include the Protocerebral Posterior Lateral 1 (PPL1) and Protocerebral Anterior Medial (PAM) clusters (**Fig 3A**) (Li et al., 2020). Previous studies have indicated that PPL1 neurons primarily encode negative reinforcement signals while PAM neurons encode positive reinforcement signals in diverse contexts, including courtship (Keleman et al., 2012; Zhao et al, 2018; Shen et al., 2023, Kim et al., 2018). Recently, it has been shown that specific dopaminergic neurons also directly regulate the behavior of flies in the social space preference paradigm (**Fig. 1A-B**) (Robinson et al., 2026).

Therefore, we used the split-GAL4 system (Aso et al., 2014) to examine the effects of *eIF4H1* knockout on the morphology of one specific neuronal subtype within each cluster. To test our hypothesis, we chose to utilize highly specific split-GAL4 drivers that label dopaminergic neurons in the Protocerebral Posterior Lateral 1 (PPL1-03) and Protocerebral Anterior Medial 10 (PAM10) clusters, which exhibit robust stereotypic anatomy and have well-defined behavioral and cognitive functions (Liu et al., 2012; Aso et al., 2010). As previously reported, we found that the PPL1-03-specific split-GAL4 labeled neurons that innervate the γ2 and α′1 lobes of the mushroom body (MB) (Li et al., 2020). In contrast, *eIF4H1*^null^ animals often lacked a robust innervation of the relevant MB lobes, and the locations of labeled neuronal soma were different from the stereotypical wild type location of PPL1-03 soma in about ∼42% (6/14) of *eIF4H1^null^* brains (**Fig. 4B-D**). Notably, we also found that *eIF4H1^null^*brains showed a significant increase in GFP-labeled ectopic neuronal soma in the suboesophageal zone (SEZ) (**Fig 4E**). Conversely, we found that the *eIF4H1^null^* allele had a relatively minor effect on the organization of neuronal in the PAM10 cluster, a subcluster of 4-5 neurons per hemisphere that innervates the β1 lobe of the MB (Li et al., 2020). We observed a mild decrease in the number of detectable PAM10 soma in *eIF4H1^null^*, and did not see a significant difference in the number of SEZ-localized ectopic cell soma in this genotype (**Fig. 4F-I**). These results indicate that the role of *eIF4H1* in driving neuronal differentiation is variable across different dopaminergic cell lineages.

**Figure 4.**
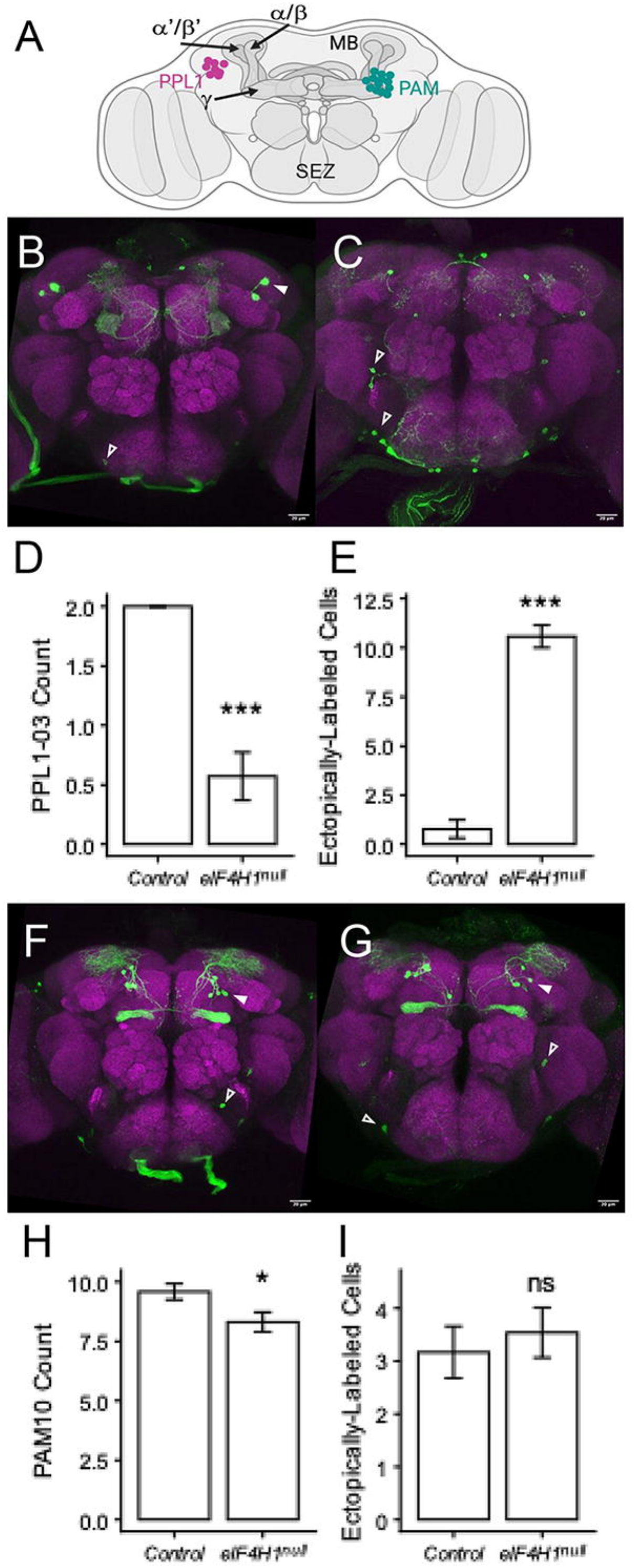
Differential effects on DAN neuron organization in *eIF4H1^null^*. A) Diagram of PPL1 and PAM neurons, with general location of respective cell soma indicated. Mushroom body (MB) and corresponding lobes indicated with arrows. SEZ corresponds to suboesophageal zone. B & C) Representative images of PPL1-03-GAL4>*UAS-mCD8::GFP* neurons in control (B) and *eIF4H1^null^*(C). Neurons labelled in green, synaptic staining (nc82) in purple for reference. Filled arrowheads indicate quantified focal cell(s) and empty arrowheads indicate representative ectopic cells. D & E) Quantification of identifiable PPL1-03 cell count and SEZ-localized ectopic cell counts in control and *eIF4H1^null^* (n = 9-14). F & G) Representative images of PAM10-GAL4>*UAS-mCD8::GFP* neurons in control (F) and *eIF4H1^null^*(G). Channel colors and arrowheads as previous. H & I) Quantification of identifiable PAM10 cell count and SEZ-localized ectopic cell counts in control and *eIF4H1^null^* (n = 12-13) Cell counts were tested with two-tailed t-test. * = *p*-value < 0.01, *** = *p*-value < 0.001, ns = not significant. Data shown as mean ± SEM.

### *The eIF4H1^null^ allele is associated with abnormal dendritic morphology of larval* sensory neurons

In addition to their characteristic sociocognitive profile, individuals with WS often exhibit abnormal sensory-motor functions, which suggests that this syndrome affects both central and peripheral neuronal pathways (Butti et al., 2024; Cheam et al., 2023; John & Mervis, 2010). This led us to investigate whether *eIF4H1* activity may also regulate aspects of sensory neuron development. The *Drosophila* larval dendritic arborization (da) neurons are periphery sensory neurons that tile that body wall, possess elaborate dendritic arbors, and are often used as a cellular model for the genetic regulation of dendritic development (Grueber et al., 2002; Emoto et al., 2006; Jan & Jan, 2010; Hatton et al., 2022; Tann et al., 2024). Notably, knockdown of members of the eIF4F complex, including eIF4A, has been shown to reduce dendritic arbor complexity in the class IV ddaC subtype (Olesnicky 2014). Therefore, we hypothesized that disruption of *eIF4H1* would lead to aberrant morphology of these cells.

We found that in comparison to wild type larvae, the *eIF4H1^null^*allele had no effect on the overall number of neuronal soma labelled by the ddaC-specific *ppk*-GAL4 driver (Ainsley et al., 2003) (**Fig. 5A,B**). However, we did find that the *eIF4H1^null^*allele is associated with reduced total dendrite length and branch numbers relative to wild type animals (**Fig. 5C-F**), but has no impact on branching frequency (**Fig 5G**) (Olesnicky et al., 2014; Brown et al., 2017). These results suggest that the activity of *eIF4H1* is necessary for driving some morphological aspects of dendritic field morphogenesis in ddaC neurons, and is likely required for the differentiation of some neuronal populations in the central and peripheral nervous systems.

**Figure 5.**
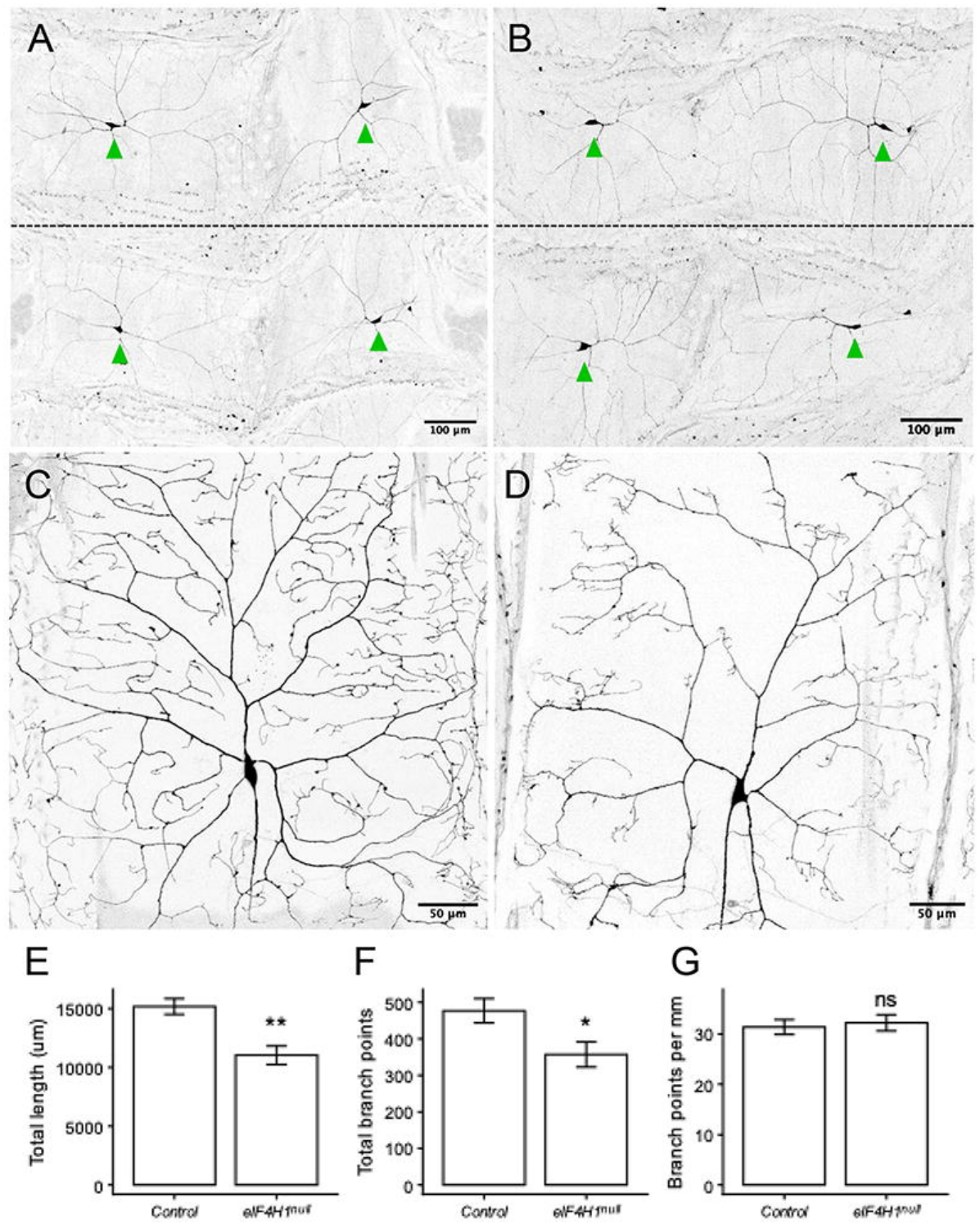
Reduced dendritic arbor size in *eIF4H1^null^*. A & B) Representative images of two segments each from Canton-S control (A) and *eIF4H1^null^*(B) larval body walls of *ppk-*GAL4>*UAS-mCD8::GFP*. Soma of each ddaC indicated with green arrowheads, dashed line indicates approximate segmental boundary. For each genotype, five different larvae were scored by counting the total number of ddaC soma across three segments. In both control and *eIF4H1^null^*, the observed count in all larvae matched the expected value of 6 cells (±0, Fisher’s exact *p-*value = 1). C & D) Representative images of ddaC neuron dendritic arbors in Canton-S control (C) and *eIF4H1^null^* (D). E-G) Quantification of dendritic arbor morphology, as measured by total dendrite length (E), total branch points (F), and branching density as represented by branch points per mm length (G) (n = 5 neurons, each from different larva). Morphology metrics were tested with two-tailed t-test. * = *p*-value < 0.05, ** = *p*-value < 0.01, ns = not significant. Data shown as mean ± SEM.

## Discussion

In this study, we identified *eIF4H1*, a component of the highly conserved eIF4F eukaryotic translation initiation complex, as a regulator of *Drosophila* social behavior and neuronal development. We find that *eIF4H1* is broadly expressed in neurons, and that both neuronal knockdown and whole-animal gene disruption causes behavioral phenotypes in multiple paradigms of fly social interaction. We show that these behavioral outcomes may be mediated, at least in part, via the selective action of eIF4H1 in some central dopaminergic neuronal populations, as well as sensory pathways. However, to our knowledge, the direct impact of eIF4H hemizygosity on sociality in humans or mammalian models has not been directly explored. Nevertheless, at least one mouse study found growth retardation, microcephaly, reduced neuron number and complexity, and severe impairments of fear-related associative learning and memory in a mouse eIF4H knockout mice (Capossela et al. 2021). Therefore, although suggestive, whether the action of eIF4H in the nervous system represents an example of a conserved social genetic toolkit remains speculative (Rittschof & Robinson, 2016; Toth & Robinson, 2007; Robinson et al., 2008).

Nonetheless, our data indicate that the eIF4H-dependent plasticity of the eIF4F complex does play an important, potentially conserved role in the regulation of the neuronal translatome. Specifically, the broader genetic toolkit framework stipulates that highly conserved gene-networks are repeatedly co-opted to drive species-specific traits by specific key regulatory nodes (*e.g.*, transcription factors) that affect the activity of numerous downstream effectors with disparate functions (Carroll 2010). Our data suggest that eIF4H1 represents one such node within critical neuronal gene networks across flies and mammals. Consequently, as a translation initiation factor, *eIF4H1* may coordinate diverse neurobiological processes through the regulation of broad sets of gene products, such as ion channels and neurotransmitters. As a node of a molecular regulatory network, this makes *eIF4H1* a conceptually “evolvable” building block that could drive evolution of the social brain at the cellular and neuronal circuit levels. The importance of the eIF4F complex in regulating the neuronal translatome is further supported by recently identified rare variants of *eIF4A2* that cause a neurodevelopmental disorder associated with intellectual disability (Paul et al., 2023). Additionally, dysregulation of eIF4E – including rare gain-of-function mutations – has been linked to autism spectrum disorder (Gkogkas et al., 2013; Huang et al., 2026).

In the context of higher brain functions associated with cognition and sociality, our data indicate that *eIF4H1* plays a specific and contextual role in shaping specific neuromodulatory pathways. Importantly, the data presented here do not directly link the observed eIF4H1-dependent changes in dopaminergic signaling to specific behavioral outcomes. However, that the dopaminergic clusters we investigated have discrete and context-specific behavioral roles is aligned with our overarching hypothesis that *eIF4H1* is not a generic housekeeping gene, but rather a tool used by the nervous system to refine the functions of specific neuromodulatory populations that play important conserved roles in cognition across insects and mammals (Barron et al., 2010; Gunaydin et al., 2014; Zhang et al., 2016; Aso & Rubin, 2016).

Although the specific mechanisms by which *eIF4H1* regulates neuronal functions and behavior remain mostly unknown, our results suggest that the regulation of the translation of some mRNAs by the eIF4F translation initiation complex can drive specific aspects of neuronal differentiation and physiology that are relevant to how nervous systems process and integrate social signals. This interpretation is well-aligned with the emerging importance of the precise spatial and temporal regulation of protein translation in neurons (Santini et al., 2013; Costa-Mattioli et al., 2009; Holt et al., 2019; Hafner et al., 2019; Van Oostrum et al., 2023; Yan et al., 2023; Kraushar, 2026). Specifically, the activity of the eIF4F complex has been shown to be critical for a variety of neuronal processes. For one, eIF4F proteins are present at the synapse and dendrite, and have been shown to be stimulated upon synaptic activity, implying that eIF4F-dependent translation control is important for synaptic plasticity, and learning and memory at the physiological timescale (Hoeffer et al., 2013; Glock et al., 2021; Eom et al., 2014). eIF4F activity has been also linked to multiple neurodevelopmental processes, including axon guidance and neural stem cell differentiation (Campbell & Holt, 2001; Lin et al., 2016; Rodrigue et al., 2025).

Our findings about the specific role of *eIF4H1* in neuronal differentiation and behavior provide one potential mechanism for how the eIF4F complex fine-tunes the neuronal translatome. Previous studies indicate that cap-dependent translation initiation by the eIF4F complex is required for the recruitment of the 43S preinitiation complex, which scans the 5′ UTR of mRNAs for the start codon. For mRNAs with long, GC-rich, G-quadruplex-containing 5′ UTRs, such as those often transcribed by neurons (Furlanis et al., 2019; Popovitchenko et al., 2020), the UTR scanning requires active unwinding of secondary RNA structures by the DEAD-box helicase eIF4A (Rogers et al., 2002; Parsyan et al., 2011; Brito Querido et al., 2024). Because of the relatively weak helicase activity of eIF4A, initiation of mRNAs with complex 5’UTRs further depends on the mutually exclusive recruitment of either the eIF4B or eIF4H RNA-binding accessory factors (Rogers et al., 2001; Özeş et al., 2011; Rozovsky et al., 2008; Sun et al., 2012; García-García et al., 2015). While both eIF4H and eIF4B paralogs are highly conserved throughout eukaryotic evolution (Sonenberg & Hinnebusch, 2009), their RNA-binding domains are different, suggesting potential mRNA target specificity (Maris et al., 2005; Cléry et al., 2008). In addition, eIF4B possesses protein-binding domains that enable protein interactions that eIF4H1 does not possess, such as with eIF4G and PABP (Özeş et al., 2011; Marintchev et al., 2009; Quintas et al., 2024). Thus, the molecular division of labor between eIF4H1 and eIF4B may represent a mechanism by which neurons fine-tune eIF4F-dependent regulation of the translatome, which could lead to the behavioral and neuroanatomical phenotypes we observed here.

In conclusion, we identified *eIF4H1* as a novel candidate sociality toolkit gene, and find that it is associated with impaired social behaviors, dopaminergic neuron organization, and sensory dendritic complexity in *Drosophila*. These results highlight the importance of the eukaryotic translation initiation complex in regulating nervous system development and function and imply that *eIF4H1* may contribute to the sociocognitive profile of individuals with Williams syndrome.

## Acknowledgements

We thank the Bloomington Drosophila Stock Center (NIH P40OD018537) for providing fly stocks. The research was supported by a Howard A. Schneiderman Graduate Fellowship from Washington University to EN, and grants NS089834 and ES025991 from the NIH, and 1545778, 1707221, and 2319060 from the NIH to YB-S.

---

**Figure S1. Reduced viability and delayed development in *eIF4H1^null^*.** A) Time course of male eclosion, data points are averages across 9 replicates. B) Fraction of total eclosed males that were *eIF4H1^null^,* each point represents a replicate, solid line indicates average fraction of 0.17, significantly lower than the Mendelian value of 0.5 (dashed line, Wilcoxon rank sum test). C) Mean eclosion day for all replicates, black dot represents average weighted CS eclosion day for given replicate, connected red dot indicates same value of *eIF4H1^null^* males from that replicate. *eIF4H1^null^* males showed an average delay of ∼1.5 days (solid lines, paired Wilcoxon rank sum test). ***, *p*-value < 0.001.

## Materials and Methods

### Fly lines used

All stocks were grown on standard food (Archon Scientific) at 25°C on a 12 hour L/D cycle. The following Bloomington Stock Center lines were used in this study, with stock numbers in parentheses. In the initial neuronal Williams ortholog screen, the pan-neuronal driver *elav-*GAL4*;UAS-Dicer2* (#25750) was crossed to the following UAS-RNAi TRIP lines (Perkins et al. 2015): *CG10903* (#57481), *eIF4H1* (#57306), *fz3* (#66951), *fz4* (#64990), *LIMK1* (#26294), *Mondo* (#27059), *Nsun5* (#32440), *Nsun5* #2 (#32980), *Syx1A* (#25811), and *Vps37B* (#44010). The TRiP *UAS-GFP* (#35786) was used as a negative control. DAN neurons were labelled with the following split-GAL4 drivers: MB296B (#68308) for PPL1-03 and MB063B (#68248) for PAM10. Larval ddaC neurons were labelled with the ppk-GAL4 (#32079) driver. To visualize DAN and larval ddaC neurons, we labelled cell membranes with *UAS-mCD8::GFP* reporter transgenes on chromosome 2 (#32186) or 3 (#32195).

### Social space assay

The general social space assay is as follows. 3-5 day old males of a given genotype were co-housed with females until 24 hours before test, at which point they were anesthetized with CO_2_ and sorted into groups for testing. On the day of the test, the flies were briefly anesthetized with ice (<30 seconds) to facilitate rapid loading into the triangular arena. The arena was a right triangle with side dimension of 16.5 cm. Flies were allowed to recover for 15 minutes, then the arena was placed upright and the flies were gently tapped to the lower left corner. After 15 minutes, a picture was taken to record the position of the flies. A custom Python-based script was used to measure the distance of each fly to the apex of the arena. In all cases, this analysis was done blind to the genotype. For a given replicate, any flies <13 cm from the arena apex were removed, as these were fallen flies not representative of the social space behavior. The distances of each remaining fly was averaged to generate the replicate’s summary statistic. All tests were performed at 25°C at ZT 5-8. Locomotion control assays were performed on the *eIF4H1-*RNAi and control genotypes used in the social space assay. Groups of ten 3-5 day old males were loaded into a cylindrical arena (diameter 2 cm) and gently tapped to the bottom. The number of males able to climb past 10 cm in 30 seconds was counted. In the WS ortholog RNAi screen, Kruskal-Wallis test was first performed to determine if any significant differences existed in the dataset. Dunn tests were then performed to test specific comparisons e.g. *eIF4H1*-RNAi to control.

### Male courtship assay

To quantify male courtship behavior, newly-eclosed males were collected and socially isolated for 3-4 days. Age-matched Canton-S virgin females were stored in groups of 5. On the day of testing, gentle mouth aspiration was used to load males and females into the arena. Flies were allowed to acclimate to the arena for 10 min, then males and females were brought together and a video was recorded for 10 min. Tests were performed at 25°C and ZT 1-4. For each trial, genotypes were blinded and courtship bouts were scored manually. Courtship latency was calculated as time to courtship onset. Courtship index = (Total time spent courting + copulating) / (Assay length – Latency). Courtship and copulation incidence metrics represent the occurrence of either event at any point in a given trial. Wilcoxon rank sum test was used to test comparisons of both metrics.

### Molecular biology and generation of new lines

The C-terminus GFP-tagged allele of *eIF4H1* was generated via CRISPR/Cas9-dependent editing by using a modified “scarless” strategy (www.flyCRISPR.molbio.wisc.edu) (Bier et al., 2016). A single *eIF4H1-*targeting sgRNA (GGTCTCTGCCACCGCGTTAA) was designed using flyCRISPR.org/target-finder to generate a double-stranded break in the last exon of *eIF4H1.* Complementary oligos for the sgRNA (IDT) were annealed and cloned into the pDCC6 plasmid (Addgene plasmid #59985), which includes the coding sequence for Cas9 (Gokcezade et al., 2014), by using the BpiI restriction enzyme (Thermo). The donor plasmid for homologous recombination was constructed by using a Golden Gate assembly (Engler et al., 2008) to recombine four DNA elements: 1) A backbone based on the pBS-GGAC-ATGC plasmid (Addgene #60949, Zhang et al, 2014) that was modified to include kanamycin resistance and a *lacZ* ORF between the BsmbI sites, enabling blue-white screening for successful donor plasmid assembly (gift from Dianne Duncan); 2) Eye-specific DsRed reporter driven by the 3XP3 promoter, flanked by two PiggyBac transposase recognition sites, which was PCR-amplified from the pHD-ScarlessDsRed plasmid (see Hill et al., 2019); 3–4) The left and right homology arms, which consisted of ∼500 bp genomic DNA fragments upstream and downstream of the sgRNA site that were synthesized as DNA gBlocks (IDT). The homology arms split the PAM sequence from the rest of the sgRNA recognition site, preventing Cas9-dependent cutting of the donor plasmid. The right homology arm included a GFP coding sequence in frame with the 3’ end of the last coding exon of *eIF4H1*, immediately upstream of the endogenous stop codon. The sgRNA-containing pDCC6 plasmids (100 ng/μL) and the donor plasmid (500 ng/μL) were co-injected into *y^-^w^-^* background. Correct genomic integration of the GFP tag was verified by screening for DsRed-positive animals, followed by sequencing of genomic PCR fragments. The final tagged *eIF4H1^GFP^*allele was generated by removing the DsRed cassette via the introduction of the piggyBac transposase (Bloomington #8285).

In parallel with the GFP-tagged *eIF4H1* allele, we also generated a C-terminal *eIF4H1*-V5 tag line. The approach and reagents for this allele were exactly as above, with the GFP sequence in the right homology arm swapped with a V5 coding sequence. We successful generated the DsRed-marked insertion and confirmed correction insertion location via genomic PCR. We subsequently used this line, without removing the DsRed cassette via piggyBac transposase, as the *eIF4H1* loss-of-function allele (*eIF4H1^null^*).

### Brain immunofluorescence and imaging

Adult brains were dissected in cold Schneider’s Medium (Thermo), fixed in 4% PFA (EMS) for 20 min at room temperature, washed with PBS containing 0.1% Triton X-100 (Sigma-Aldrich), and blocked overnight at 4°C in SuperBlock (Thermo). Brains were then incubated in primary antibody mix overnight at 4°C, washed thoroughly with 0.1% PBS-T, and incubated in secondary antibody mix overnight at 4°C. All antibodies were diluted in SuperBlock, see below sections for specific mix compositions. Following final washes in 0.1% PBS-T, brains were mounted onto slides using Vectashield Vibrance mounting medium (Vector Laboratories). Size 2 cover slips were used as “spacers” to prevent compression of brains. Brains were imaged using a laser scanning confocal microscope (Leica Sp8 Lightning). All confocal images were analyzed and processed in Fiji (Schindelin et al., 2012).

### eIF4H1^GFP^ staining

To stain for eIF4H1^GFP^, brains of 1-5 day old *eIF4H1^GFP^/FM6* females were dissected and stained with primary antibodies rabbit anti-GFP (ThermoFisher, A-11122) at 1:1000 and rat anti-ELAV (Developmental Studies Hybridoma Bank, 7E8A10) at 1:200. Secondary antibodies in this staining were Alexa Flour 488-conjugated donkey anti-rabbit (ThermoFisher, A-21206) and Alexa Flour 568-conjugated goat anti-rat (ThermoFisher, A-11077), each diluted 1:2000. Whole brain images were taken with a 20X objective, and regions of interest were imaged with a 40X objective.

### *eIF4H1^null^* viability and behavioral experiments

We first moved the *eIF4H1^null^* allele to our standard Canton-S (CS) background. Heterozygous *eIF4H1^null^* females were crossed to CS males and the 3XP3>DsRed marker was used to select the null allele in the next generation. This process was repeated for 6 generations, with new CS males being used every 3-4 generations subsequently throughout the maintenance of the stock. To quantify the effects of the *eIF4H1^null^*allele on viability and developmental timing, nine crosses were established containing ten mated heterozygous (*eIF4H1^null^* /+) females each. Females were allowed to lay eggs in fresh vials for three consecutive 24-hour intervals, generating three egg-lay cohorts per cross. The number of *eIF4H1^null^*and CS males eclosing from each cohort was recorded daily at ZT3 for consistency. For each cross replicate, male progeny counts were summed across the three cohorts to calculate the fraction of eclosed males carrying the *eIF4H1^null^*allele. Mean eclosion day was calculated separately for *eIF4H1^null^*and CS males as the weighted average day of adult emergence. Statistical analyses were performed using cross-level summary values (n = 9 biological replicates). *eIF4H1^null^* male recovery was compared to the expected Mendelian frequency of 0.5 using a one-sample Wilcoxon signed-rank test, while differences in mean eclosion day were tested using a paired Wilcoxon signed-rank test. To obtain males for subsequent behavioral testing, groups of heterozygous *eIFH1^null^* females were crossed to CS males. Resulting male progeny were either CS (used as controls) or *eIFH1^null^*(experimental group). Male courtship assays were performed on these *eIF4H1^null^*and CS males as detailed above. Social space assays were performed also as above, but with groups of 15 males being used in each test due to the lower viability of *eIFH1^null^* males. Wilcoxon rank sum tests were used to determine statistical significance.

### DAN neuron imaging and quantification

To visualize DAN neurons in the background of *eIF4H1^null^*, an *eIF4H1^null^/FM6; UAS-mCD8::GFP*/CyO line was generated via standard genetic crosses. Experimental animals were obtained by crossing DAN split-GAL4 drivers to this line, while control animals were generated using a corresponding *+;UAS-mCD8::GFP* line. Progeny carrying the *UAS-GFP* and both split-GAL4 elements were selected for dissection and imaging. To image PPL1-03 and PAM10 neurons, 1-5 day old males of the appropriate genotypes were dissected and stained with primary antibodies rabbit anti-GFP (as above) and mouse anti-*brp* (nc82, Developmental Studies Hybridoma Bank) at 1:50. Secondaries antibodies were Alexa Flour 488-conjugated donkey anti-rabbit and Alexa Flour 647-conjugated donkey anti-mouse (ThermoFisher, A-31571), each diluted 1:2000. To quantify the effects of *eIF4H1^null^* on PPL1-03 and PAM10 organization, brains were dissected and stained as above, then mounted in batches of 5 brains per slide. Counts of identifiable focal (PPL1-03 or PAM10) and ectopic cells were scored manually, blind to genotype. Focal cells were identified based on cell soma position and stereotypical mushroom body innervation pattern. Ectopic cells were defined as those localized to the general area of the suboesophageal zone (SEZ). We note that the PPL1-03 split-GAL4 driver also occasionally labelled PPL1-05, which was easily distinguishable based on alternative morphology and was not included in either focal or ectopic cell counts. Two-tailed t-tests were used to compare metrics across genotypes.

### ddaC neuron imaging and quantification

To visualize larval ddaC neurons, we generated a recombined *ppk-GAL4>UAS-mCD8::GFP* transgene on chromosome 3 through standard genetic crossing. This line was crossed to heterozygous *eIF4H1^null^/+* females, and male 3^rd^ instar larva of the appropriate genotype were collected for imaging. To count the number of ddaC neurons per segment across the larval body wall, larvae were dissected in ice-cold Schneider’s Medium, filleted with the dorsal interior body wall facing up, and pinned to flatten the body wall. Larvae were fixed and washed with the same procedure and reagents as brains, then the pins were removed and larval tissues moved to 1.5 mL tubes for the remainder of the staining. The primary antibody used was the same rabbit anti-GFP as above at 1:1000, followed by the Alexa Flour 488-conjugated donkey anti-rabbit secondary at 1:2000. After the completion of staining, body walls were mounted in 70% glycerol (Sigma-Aldrich) and imaged via laser scanning confocal microscopy. While useful for imaging entire larval body walls, we found that fixation and staining was not amenable for accurate representation of individual ddaC dendritic arbors due to distortion and folding of the body wall after dissection. Therefore, to quantify ddaC dendritic morphology, we performed live imaging on larvae of the same genotypes as above. Larvae were first anesthetized briefly with diethyl ether (Sigma-Aldrich), then mounted in 70% glycerol. Several strips of double-sided tape were used to ensure that larvae were not crushed. Images of ddaC dendritic arbors were captured within 30 minutes of mounting via laser scanning confocal microscopy. Resulting ddaC dendritic arbors were traced and quantified with the SNT program in FIJI (Arshadi et al., 2021). Two-tailed t-tests were used to compare morphological metrics between control and *eIF4H1^null^*.

## Statistical analysis

All indicated statistical comparisons were performed with custom R scripts (version 4.6.0, 2026) in the RStudio program (Posit, 2026).

