## Supplementary figures and images for "Regulation of sociality and neuronal development by the translation initiation factor *eIF4H1* in *Drosophila*"

### Supp Fig. 1

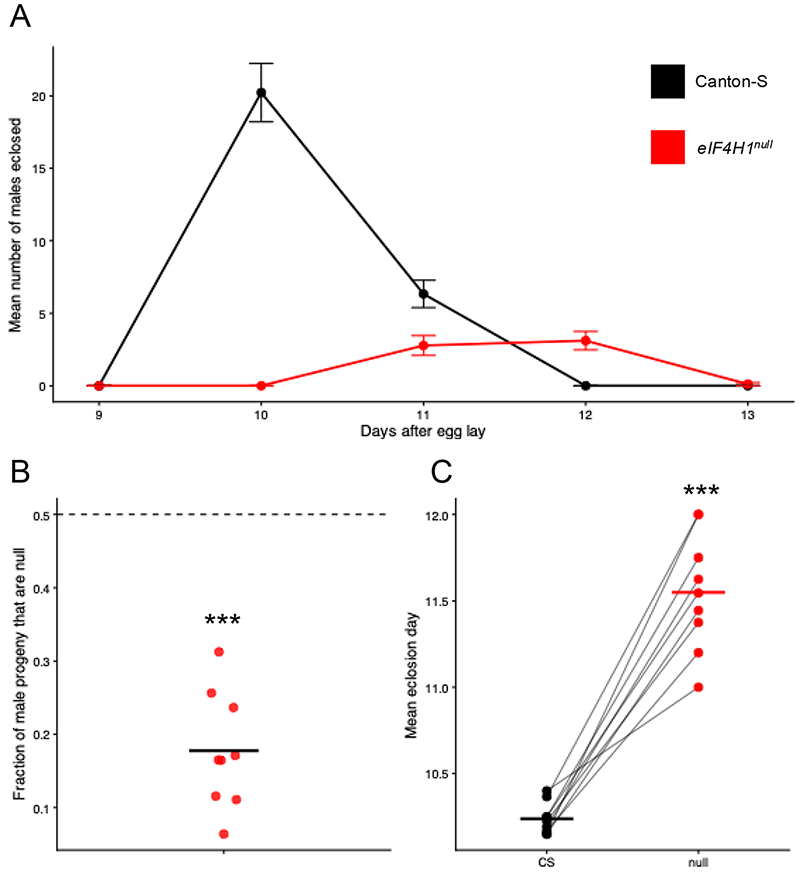
